# The *Streptococcus pyogenes* hyaluronic acid capsule promotes experimental nasal and skin infection by preventing neutrophil-mediated clearance

**DOI:** 10.1101/2022.04.07.487539

**Authors:** Jacklyn R. Hurst, Blake A. Shannon, Heather C. Craig, Stephen W. Tuffs, John K. McCormick

## Abstract

*Streptococcus pyogenes* is a globally prominent human-specific pathogen responsible for an enormous burden of human illnesses, including >600 million pharyngeal and >100 million skin infections each year. Despite intensive efforts that focus on invasive indications, much remains unknown about this bacterium in its natural state during colonization of the nasopharynx and skin. Using acute experimental infection models in HLA-transgenic mice, we evaluated how the hyaluronic acid (HA) capsule contributes to *S. pyogenes* MGAS8232 infection within these limited biological niches. Herein, we demonstrate that HA capsule expression promotes bacterial burden in murine nasal turbinates and skin lesions by resisting neutrophil-mediated killing. HA capsule production is encoded by the *hasABC* operon and compared to wildtype *S. pyogenes* infections, mice infected with a Δ*hasA* mutant exhibited over a 1000-fold CFU reduction at 48-hours post­nasal challenge, and a 10,000-fold CFU reduction from skin lesions 72-hours post-skin challenge. HA capsule expression contributed substantially to skin lesion size development following subdermal inoculations. In the absence of capsule expression, *S. pyogenes* revealed drastically impeded growth in whole human blood and increased susceptibility to killing by isolated neutrophils *ex vivo*, highlighting its important role in resisting phagocytosis. Furthermore, we establish that neutrophil depletion in mice, but not macrophage depletion, recovered the reduced burden by the Δ*hasA* mutant in both the nasopharynx and skin. Together, this work confirms that the HA capsule is a key virulence determinant during acute infections by *S. pyogenes* and demonstrates that its predominant function is to protect *S. pyogenes* against neutrophil-mediated killing.

**AUTHOR SUMMARY:** *Streptococcus pyogenes* is a globally disseminated and human-adapted bacterial pathogen that has evolved an arsenal of evasion strategies to overcome and escape host immune clearing mechanisms. Many strains of *S. pyogenes* are covered by a polysaccharide capsule composed of hyaluronic acid (HA) that is widely recognized to promote severe infections. In this study, we demonstrate using the encapsulated *S. pyogenes* MGAS8232 strain that the HA capsule is a key virulence factor that facilitates non-invasive infections of the nasopharynx and skin. Although bacterial adhesion and entry into host cells was impeded by HA capsule expression, we show that the key function for both nasal and skin infections is to protect *S. pyogenes* from neutrophil-mediated killing. Depletion of neutrophils, but not macrophages, recovered the low bacterial burden by unencapsulated *S. pyogenes* at both sites of infection. Our findings outline an important interaction between the HA capsule and neutrophils in the establishment of acute upper respiratory and skin infections by *S. pyogenes*.

## INTRODUCTION

*Streptococcus pyogenes* (the group A *Streptococcus*) is a globally prominent, human-adapted bacterial pathogen responsible for an enormous burden of disease [1]. While *S. pyogenes* exists primarily as an asymptomatic commensal in up to 12% of school-aged children [2], more than 600 million cases of pharyngitis and 100 million cases of skin infections are recorded each year, and at least 18.1 million people worldwide currently suffer from serious post-infection sequelae resulting in over 500,000 annual deaths [1].

Many strains of *S. pyogenes* produce a high molecular weight hyaluronic acid (HA) polysaccharide capsule that presents distinct mucoid colony morphology when grown on solid media. The HA capsule is composed of repeating disaccharide units of glucuronic acid and *N*-acetylglucosamine (GlcNAc) and is structurally identical to HA found within the human extracellular matrix (ECM), and therefore, is immunologically inert [3]. Capsule production is encoded by the *hasABC* genetic locus involved in HA biosynthesis [4–6]. The first gene in the operon, *has*A, encodes hyaluronate synthase [7, 8]; the second gene, *hasB*, encodes UDP-glucose 6-dehydrogenase [9]; and the third gene, *hasC,* encodes UDP-glucose pyrophosphorylase [10]. Although proteins encoded by *hasB* and *hasC* are enzymatically active, they are not individually essential for HA capsule synthesis [11, 12]. Interestingly, expression of the *hasA* gene is the only fundamental gene required for the production of the HA polymer from UDP-glucuronic and UDP-GlcNAc sugar precursors [7]. The amount of capsule produced can vary widely among individual strains, regulated by growth conditions and in response to changes in the host environment. Maximal HA capsule production occurs during early and mid-exponential phase *in vitro*, followed by capsule shedding during stationary phase [5]. These observations are further supported *in vivo* as introduction of *S. pyogenes* into the pharynx of non-human primates or into the mouse peritoneum induces high levels of HA capsule gene transcription within 1-2 hours of inoculation [13], suggesting that capsule expression has an important function during initial stages of colonization.

Early pioneering studies using encapsulated M18 and M24 serotypes revealed that transposon mutants lacking the HA capsule had ∼100-fold increases in the LD50 using invasive intraperitoneal infections in CD1 mice [14–16], and further discovered that capsule expression was strongly selected for during pharyngeal colonization of BALB/c mice [15]. Following intratracheal inoculation of the mouse-adapted B514 *S. pyogenes* strain, the HA capsule promoted chronic throat colonization, pneumonia, and secondary systemic infections in C57BL/10SnJ mice [17]. Furthermore, encapsulation was shown to enhance persistent colonization of *S. pyogenes* in the pharynx of baboons as unencapsulated mutants were cleared more quickly [18]. Together, encapsulation appears to offer *S. pyogenes* a powerful survival advantage for colonization and dissemination.

Prior studies have shown that the HA capsule binds to the cell surface ligand CD44 to mediate adherence to epithelium [19, 20], which can induce host cell signaling events that disrupts tight junctions to promote invasion [21]. Furthermore, the HA capsule can also specifically bind to <u>ly</u>mphatic <u>v</u>essel <u>e</u>ndothelial receptor-1 (LYVE-1) expressed in lymph node sinuses and lymphatic vessles, to promote dissemination to draining lymphnodes via the lymphatic system using a mouse thigh muscle infection model [22]. However, reduced binding efficiencies by encapsulated strains have also been observed [19, 23]. Removal of the capsule by genetic inactivation of the *has* operon can also promote robust invasion of cultured epithelial cells, although once internalized, *S. pyogenes* is rapidly killed [24]. By producing a molecule ubiquitously expressed by its host, molecular mimicry enables *S. pyogenes* to avoid detection by host immune surveillance and increases resistance to phagocytic-mediated killing. In several experiments, unencapsulated mutants display significant susceptibility to complement-dependent phagocytic killing by human blood compared to their encapsulated parental strains [16,25,26].

Although nearly all *S. pyogenes* strains encode the *has* operon, the capsule is not universally present in all isolates. For example, M4 and M22 serotypes, and some M89 serotypes, do not contain the *hasABC* operon and thus cannot express HA capsule [27, 28], suggesting that capsule expression is not essential for pathogenicity across all serotypes. Furthermore, studies in human carriers and other primate models have also identified mutations that reduced or eliminated capsule production in long-term carriage isolates [29, 30]. Therefore, differential regulation of the HA capsule may consequently offer an important survival adaptation in specific host environments. Thus, while it is recognized that encapsulation may be advantageous for bacterial virulence, mechanisms whereby it promotes acute *S. pyogenes* infections *in vivo* merit further investigation [31]. In this work, we aimed to further evaluate the role of the HA capsule in two non-invasive murine infection models using a precise genetic deletion of the *hasA* gene. Herein, we demonstrate using the encapsulated *S. pyogenes* MGAS8232 strain that the HA capsule is a key virulence factor for non-invasive nasopharyngeal and skin infections. Though removal of the capsule permitted bacterial invasion into host cells, we demonstrate that the key function for both *in vivo* nasal and skin infections is to protect *S. pyogenes* from neutrophil-mediated killing.

## RESULTS

### The *S. pyogenes* HA capsule promotes nasopharyngeal infection that enhances a cytokine response that supports the recruitment of neutrophils

In order to evaluate the influence of the HA capsule during experimental infections, we used the pG^+^host5 integration plasmid [32] (**Table 1**) to generate a markerless 1,212-bp in-frame deletion of *hasA* in the M18 serotype rheumatic fever isolate *S. pyogenes* MGAS8232 [33] (**Table 1**). M18 serotypes are well known for being highly encapsulated, a phenotype that has been traced to the mutations within the RocA regulatory protein [34]. The isogenic nature of the *hasA* mutant was determined by PCR and DNA sequencing with primers that flanked the deleted *hasA* region (**Table S1**) and whole genome sequencing analysis. The correct Δ*hasA* deletion was confirmed, but compared to the wildtype MGAS8232 strain, two non-synonymous single nucleotide polymorphisms (SNPs) within *pstI* gene encoding the cytosolic protein enzyme I in the phosphoenolpyruvate phosphotransferase system (PTS) were also identified and resulted in two amino acid substitutions (Leu194Phe and Ser306Phe) (**Table S2**). Thus, we generated a complementation strain using the pDC*erm* plasmid [35] (**Table S1**) by expressing the *hasA* gene and its native *hasA* promoter in the Δ*hasA* mutant background (MGAS8232 Δ*hasA* + *hasA*). As predicted, MGAS8232 Δ*hasA* lost the large mucoid colony phenotype on sheep blood agar compared to wildtype MGAS8232, and capsule production was restored in the *hasA*-complemented strain (**Figure 1A**).

**Figure 1.**
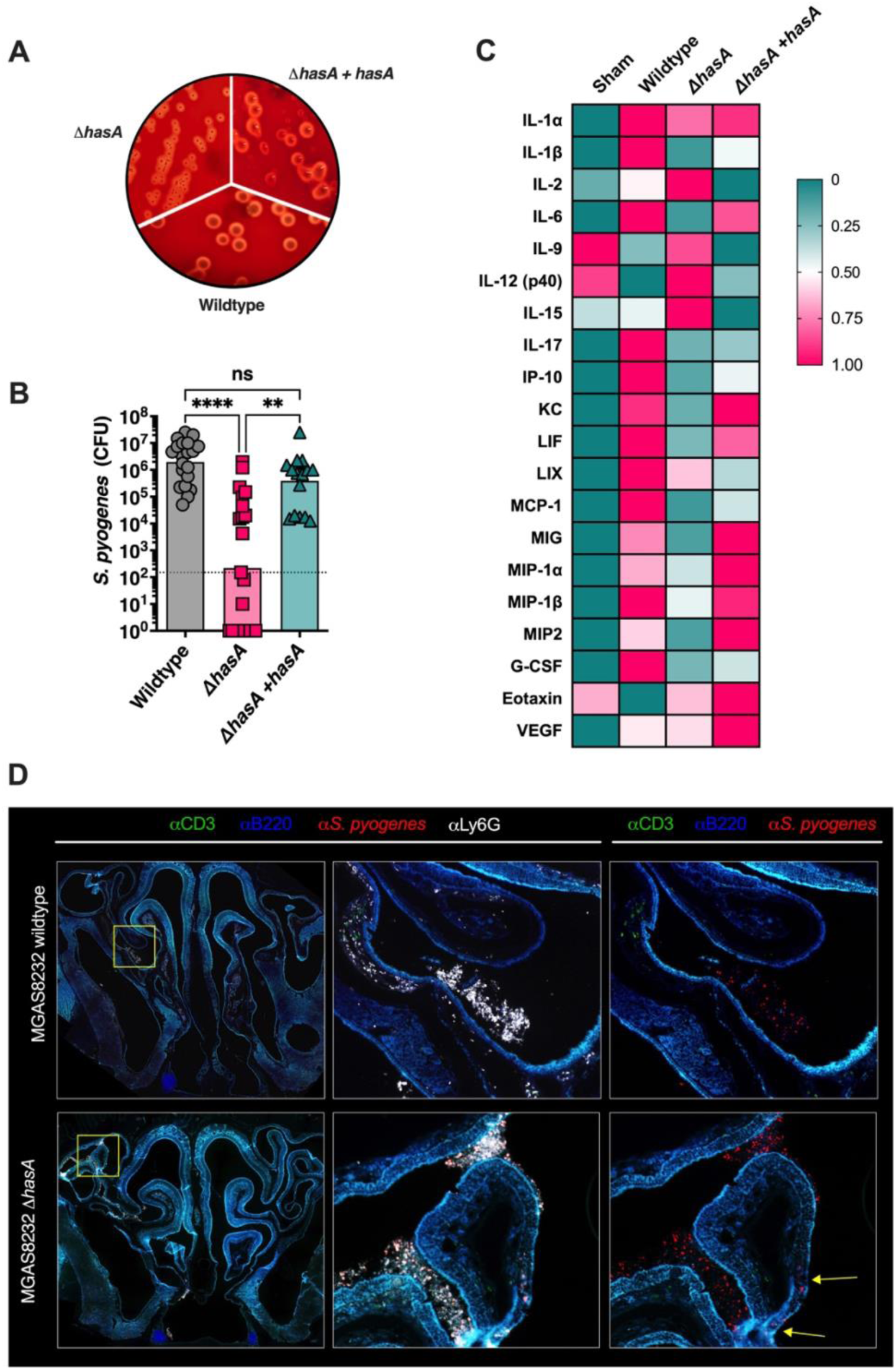
Hyaluronic acid expression by *Streptococcus pyogenes* promotes nasal infection in B6_HLA_ mice. **(A)** *S. pyogenes* constructs streaked onto TSA + 5% sheep blood agar plates **(B)** B6_HLA_ mice were administered ∼10^8^ CFUs of *S. pyogenes* MGAS8232 wildtype, Δ*hasA,* or Δ*hasA +hasA* strains intranasally and sacrificed 48 h later. Data points represent CFUs from cNTs of individual B6_HLA_ mice. Horizontal bars represent the geometric mean. Significance was determined by Kruskal Wallis one-way ANOVA with Dunn’s multiple comparisons test (****, *P* < 0.0001; **, *P* < 0.01). The horizontal dotted line indicates the theoretical limit of detection. **(C)** Heat-map of cytokine responses in cNTs of B6_HLA_ mice during *S. pyogenes* infection. Data shown represent normalized median cytokine responses from cNTs (*n* ≥ 3 per group). **(D)** Immunohistochemistry of infected cNTs at 24 h post-infection with wildtype *S. pyogenes* MGAS8232 and the Δ*hasA* mutant. Sections were stained with α-*S. pyogenes* (red), αB220 (blue), αCD3 (green), and αLy6G (white) antibodies. Panels are a close-up view from the boxed section. Arrows indicate regions with internalized *S. pyogenes*.

**Table 1.** Bacterial strains and plasmids used in this study.

| Strain/plasmid | Description | Source |
| --- | --- | --- |
| *Streptococcus pyogenes* |  |  |
| MGAS8232 | M18 serotype isolated from a patient with acute rheumatic fever (GenBank accession: NC_003485.1) | [33] |
| MGAS8232 $\Delta hasA$ | *hasA* deletion mutant derived from MGAS8232 | This study |
| MGAS8232 $\Delta hasA$ + *hasA* | MGAS8232 $\Delta hasA$ containing complementation plasmid pDCerm expressing the *hasA* gene | This study |
| *Escherichia coli* |  |  |
| XL1-Blue | General cloning strain | Stratagene |
| Plasmids |  |  |
| pG<sup>+</sup>host5 | Temperature-sensitive Gram-positive/*E. coli* shuttle vector; Erm<sup>r</sup> | [32] |
| pG<sup>+</sup>host5::$\Delta hasA$ | pG<sup>+</sup>host5 with *hasA* flanking regions inserted | This study |
| pDCerm | Erm<sup>r</sup> derivative of streptococcus-*E. coli* shuttle vector pDC123 (*erm* of Tn916$\Delta E$) replacing *cat* | [35] |
| pDCerm::*hasA* | pDCerm expressing *hasA* gene for plasmid-based complementation | This study |

Erm<sup>r</sup> - erythromycin resistance
*Cat* - chloramphenicol acetyltransferase

The human-specific tropism of *S. pyogenes* represents a major challenge when conducting experimental infection models. We previously demonstrated that the use of transgenic mice that express human MHC class II molecules (herein referred to as B6_HLA_ mice) greatly enhances *S. pyogenes* nasopharyngeal infection due to the selective specificity of superantigens for human MHC class II molecules [36]. Using this infection model, we examined the influence of HA capsule expression on bacterial burden during acute nasopharyngeal infection through nasal inoculation (∼1×10^8^ CFU) of B6_HLA_ mice using the three MGAS8232 strains. The Δ*hasA* mutant resulted in a >1000-fold reduction in bacterial CFUs from the nasal mucosa at 48 hours compared to infection by wildtype *S. pyogenes* MGAS8232 (**Figure 1B**). We confirmed that capsule expression was specifically required for the nasopharyngeal infection phenotype as the complemented strain with restored capsule expression phenocopied the wildtype infection (**Figure 1B**). Consistent with the non-invasive nature of the model [36], mean bacterial dissemination of wildtype *S. pyogenes* remained below the limit of detection in the lungs, liver, spleen, heart, and kidneys (**Figure S1**). We conclude that HA capsule expression improves experimental *S. pyogenes* nasopharyngeal infection in B6_HLA_ mice, and removal of the capsule does not increase bacterial dissemination of *S. pyogenes* MGAS8232.

To further assess the nasopharyngeal environment during wildtype *S. pyogenes* and Δ*hasA* infections, we conducted a multiplex cytokine and chemokine array using infected nasal turbinate homogenates [37]. Quantitative data is shown in supplemental figures (**Figure S2**) and summarized as a heatmap showing normalized cytokine responses for any cytokine with an average concentration above 20 pg ml^-1^ within a treatment group. (**Figure 1C**). Uninfected mice demonstrated no apparent inflammatory signature from the cNT homogenates whereas robust cytokine responses were evident in wildtype-infected mice and included: Th1-type cytokines (IL-1α and IL-1β); Th17-type cytokines (IL-6 and IL-17); chemokines (KC, IP-10, MCP-1, MIP-1α, MIP-1β, MIG, MIP-2, LIF and LIX); and growth factors (G-CSF) (**Figure 1C**, **Figure S2**). In contrast, Δ*hasA*-infected cNTs presented a reduced inflammatory signature and revealed similar cytokine expression profiles as uninfected mice. Particularly, reductions were detected with pro-inflammatory cytokines, such as IL-1β, IL-6, and IL-17, and those involved in monocyte and neutrophil recruitment, including KC, IP-10, MCP-1, MIP-1β, MIG, and G-CSF (**Figure 1C**, **Figure S2**). Restoring capsule expression in the Δ*hasA* mutant background induced a moderately inflamed environment, trending for greater concentrations of KC, MIP-1α, MIG, MIP-2, and G-CSF compared to Δ*hasA* mutant infections. Interestingly, IL-2, IL-12 (p40), IL-15, and IL-9 concentrations were higher in cNTs challenged with Δ*hasA* mutant compared to wildtype or Δ*hasA* + *hasA* strains; however, concentrations for these cytokines did not drastically differ from uninfected murine cNTs (**Figure S2**). There were also numerous cytokines that did not have average concentrations above 20 pg ml^-1^ in any treatment groups, were not different between treatment groups, or did not conform to any obvious trends (**Figure S2**). These cytokine trends suggest that at 48 hours post-nasal inoculation with *S. pyogenes,* HA capsule expression is associated with higher concentrations of cytokines and chemokines that support inflammation and monocyte and neutrophil function.

To gain further insight into the interaction between *S. pyogenes* and host immune cells during nasopharyngeal infection, MGAS8232 wildtype and Δ*hasA*-infected cNTs were harvested and cryopreserved for immunohistochemistry. Due to the possibility that Δ*hasA* mutant could be completely cleared by 48 hours post-infection (**Figure 1B**), infected cNTs were collected at 24 hours post-infection and sections were stained with α-*S. pyogenes* (red), α-B220 (blue), α-CD3 (green), and α-Ly6G (white) fluorescent antibodies where indicated. Sections revealed that *S. pyogenes* was present with robust α-Ly6G neutrophil signals in both wildtype and Δ*hasA*-infected cNTs (**Figure 1D**). By 24 hours, neutrophils infiltrated to the central areas where *S. pyogenes* resided, whether the HA capsule was expressed or not (**Figure 1D**). Although few differences in immune cell percentages have been detected within the cNTs of wildtype-infected B6_HLA_ mice by 48 hours [36], infected nasal passages have demonstrated increased trends for neutrophil populations (GR1^+^) during wildtype *S. pyogenes* MGAS8232 infection [36], entirely consistent with these immunofluorescence experiments. Notably, *S. pyogenes* Δ*hasA,* but not wildtype, was detected within the epithelial cell layer in some areas (**Figure 1D**, denoted by arrows in the right panel). Together, these observations are consistent with a model that neutrophils are recruited to murine cNTs by 24 hours post-infection with either wildtype MGAS8232 or Δ*hasA* nasopharyngeal infection, however, by 48 hours Δ*hasA* mutants are rapidly cleared and cannot remodel the nasopharynx to express favourable inflammatory responses.

### The hyaluronic acid capsule blocks *S. pyogenes* adherence and invasion of pharyngeal cells and protects against killing by neutrophils *in vitro*

If *S. pyogenes* successfully escapes mucocilliary clearance, it proceeds to target and adhere to the underlying epithelial surface. Multiple studies have described the HA capsule as an important adhesin [19, 20], and we next sought to evaluate and compare adherence capabilities of wildtype *S. pyogenes* MGAS8232 with its unencapsulated mutant. Collagen type IV and fibronectin make up a significant portion of the nasopharyngeal ECM, and therefore, bacterial binding to these structures may contribute to streptococcal infection [38–41]. Collagen type IV is the primary component of the ECM basement membrane that underlays epithelial cells, and fibronectin, while only a minor component, is frequently secreted to mediate adhesion and migration of host cells [42, 43]. Bacterial binding was assessed by inoculating *S. pyogenes* onto wells pre-coated with either collage type IV or fibronectin. A decline in both fibronectin binding and collagen type IV binding was observed for the Δ*hasA* mutant (**Figure 2A and 2B**) demonstrating that under *in vitro* growth conditions*, S. pyogenes* HA capsule can likely adhere to the ECM through interactions with both collagen type IV and fibronectin.

**Figure 2.**
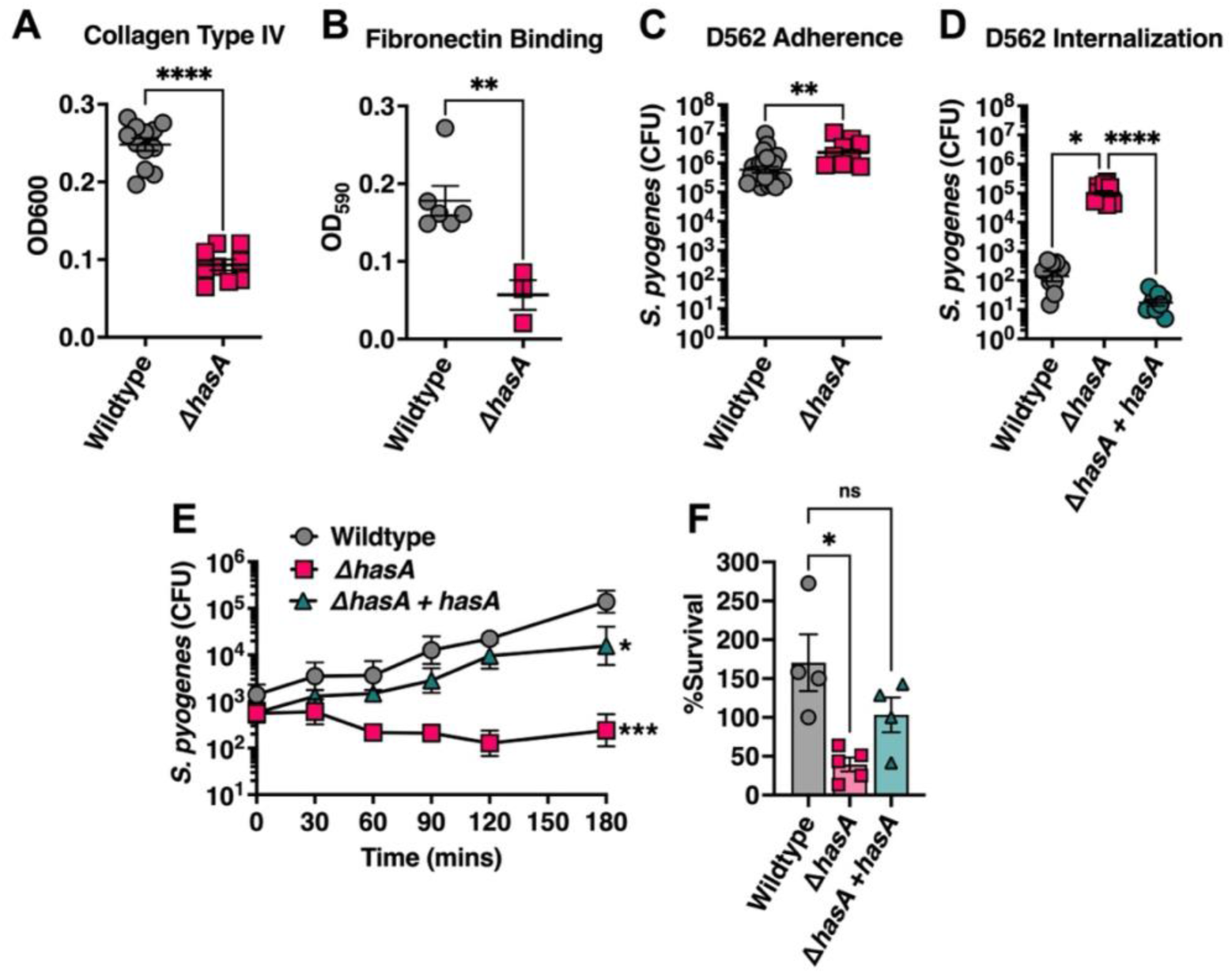
The *S. pyogenes* HA capsule inhibits host cell invasion but promotes survival from neutrophil-mediated killing. Binding of *S. pyogenes* to wells pre-coated with 1 µg of human extracellular membrane components **(A)** fibronectin and **(B)** collagen type IV **(C)** Adhesion of *S. pyogenes* to D562 pharyngeal epithelial cells. Confluent cell monolayers were cultured with *S. pyogenes* (MOI of 100) for 2 h at 37°C + 5% CO_2_. Cells were washed with PBS and lysed with Triton X-100 for enumerating remaining adherent bacteria. **(D)** Internalization of *S. pyogenes* into D562 cells. Confluent D562 cells were cultured with *S. pyogenes* (MOI of 100) for 2 h at 37°C + 5% CO_2_ followed by 1 h in media supplemented with 100μg mL^-1^ of gentamycin. Bars represent mean CFUs ± SEM (n = 3). Statistical differences were evaluated by unpaired *t*-test (**A – C**) (**, *P* < 0.01; ****, *P* < 0.0001) or (**D**) one-way ANOVA (*, *P* < 0.05; ****, *P* < 0.0001). **(E)** Whole human blood survival assay. Heparinized blood from human donors were inoculated with ∼10^3^ CFUs of *S. pyogenes* MGAS8232 at 37°C with rotation for 3 hours. Data points represent geometric mean CFUs ± SD at each timepoint. Statistical significance was determined using one-way ANOVA with Friedman test (***, *P* < 0.001). **(F)** Neutrophil survival assay. Neutrophils were isolated from human blood by density centrifugation and inoculated with opsonized *S. pyogenes* at a multiplicity of infection of 10. Surviving bacteria were enumerated after 60 mins at 37°C with rotation and calculated as the difference between survival in the no neutrophil control and in the presence of neutrophils. Each data point represents *S. pyogenes* CFUs from individual donors. Data shown are the means of percent survival ± SD. Statistical analyses were performed using one-way ANOVA with Kruskal-Wallis test (*, *P* < 0.05).

Bacterial adherence was also evaluated using the pharyngeal cell line Detroit-562 (D562) due to its similarity of surface molecules with non-transformed pharyngeal cells [43], and its ability to induce streptococcal superantigens and DNAses that are otherwise unexpressed by *S. pyogenes* when cultured on its own [44, 45]. There was a slight but statistically significant increase in the amount of *S. pyogenes* that adhered to D562 cells when HA capsule expression was absent (**Figure 2C**). Despite the ability to bind collagen type IV and fibronectin, these results conflict with reports indicating the HA capsule contributes substantially to bacterial adhesion properties, and suggests that the HA capsule may function in part to mask adhesins on the bacterial cell wall and obstruct adherence, at least with *S. pyogenes* MGAS8232 [15, 20]. Next, we aimed to characterize the capicty of these constructs for epithelial cell internalization. We found a dramatic ∼1000-fold increase in Δ*hasA* mutants recovered from lysed D562 cells following gentamycin treatment (**Figure 2D**). Given this dramatic phenotype, we also evaluated the capsule complemented strain (Δ*hasA* + *hasA*), which completely lost the invasion phenotype (**Figure 2D**). These data demonstrate that HA capsule expression by *S. pyogenes* MGAS8232 inhibits adhesion and represses internalization into pharyngeal epithelial cells.

Following attachment to epithelial cell surfaces, a critical mechanism during early colonization stages is to evade host immune responses. *S. pyogenes* MGAS8232 resistance to bacteriolysis was investigated to determine if the HA capsule improves immune evasion. Compared with wildtype *S. pyogenes* MGAS8232, growth and survival in whole human blood was markedly attenuated (∼3 logs) in the absence of capsule expression (**Figure 2E**). Upon earlier histological assessment of infected nasal turbinates, neutrophils had accumulated in regions surrounding both wildtype *S. pyogenes* and Δ*hasA* at 24 hours post-infection, yet the Δ*hasA* strain had significantly less bacterial burden by 48 hours (**Figure 1B, 1C**). Furthermore, a significant decline in cytokines and chemokines involved in recruiting, modulating, and activating neutrophils were detected by 48 hours with Δ*hasA* infection (**Figure 2B; Appendix 3**). Therefore, we next sought to examine how the HA capsule resists neutrophil activity specifically. Unencapsulated bacteria were more sensitive to neutrophil-mediated killing demonstrated by a significant decline in Δ*hasA* mutants that survived in the presence of freshly isolated human neutrophils (**Figure 2F**). Indeed, the increased susceptibility in each condition was rescued with complementation of capsule expression in the MGAS8232 Δ*hasA* + *hasA* strain (**Figure 2E and 2F**). These results confirm an important role for the capsule in promoting resistance to killing by neutrophils, and thus, may provide an innate immune role for HA capsule during early stages of acute infection.

### Depletion of neutrophils, but not macrophages, restores *S. pyogenes* Δ*hasA* bacterial load during nasopharyngeal infection

Preventing opsonophagocytic bacterial clearance is one of the main proposed mechanisms for the HA capsule and has been repeatedly investigated using various *in vitro* bacterial survival assays [16,25,26]. Since neutrophil influx is a major feature of our experimental nasopharyngeal model and during natural infections [46], we aimed to explore the importance of neutrophils during nasopharyngeal infection and determine whether preventing phagocyte-mediated killing is a key molecular process by which HA capsule functions in this model. For this purpose, mice were depleted of neutrophils by administering the αLy6G monoclonal antibody (**Figure 3A**), which effectively depletes neutrophils from the peripheral blood of mice [47], with rat IgG2a used as an isotype control. We examined the effect of depleting neutrophils at both 24-and 48-hours after nasal challenge with wildtype MGAS8232 or Δ*hasA* strains. Following the depletion of neutrophils, no differences in the amount of wildtype *S. pyogenes* recovered from infected cNTs were observed in neutrophil depleted mice compared to control mice at either 24-or 48-hours post-infection (24h, *p* = 0.5802; 48h, *p* = 0.2803), indicating that neutrophil depletion in this model does not impact nasopharyngeal infection by wildtype *S. pyogenes* (**Figure 3B**). As expected, isotype treated mice showed a reduction for the Δ*hasA* mutant at both 24 hours (*p* < 0.05) and 48 hours (*p* < 0.0001) compared to wildtype-infected control mice (**Figure 3B**). While there was no difference in recovery of the Δ*hasA* mutant between control and neutrophil depleted mice at 24 hours post-infection (*p* = 0.9808), there was a significant increase in the Δ*hasA* mutant bacterial burden at 48 hours post-infection in neutrophil depleted mice compared to control mice (*p* < 0.05). Furthermore, no statistical differences were observed between Δ*hasA* mutants recovered from neutrophil depleted mice and control mice receiving wildtype *S. pyogenes* (*p* = 0.0989). Overall, these findings suggest neutrophil-mediated clearance mechanisms contribute substantially to the lower burden of unencapsulated *S. pyogenes* in the nasopharynx.

**Figure 3.**
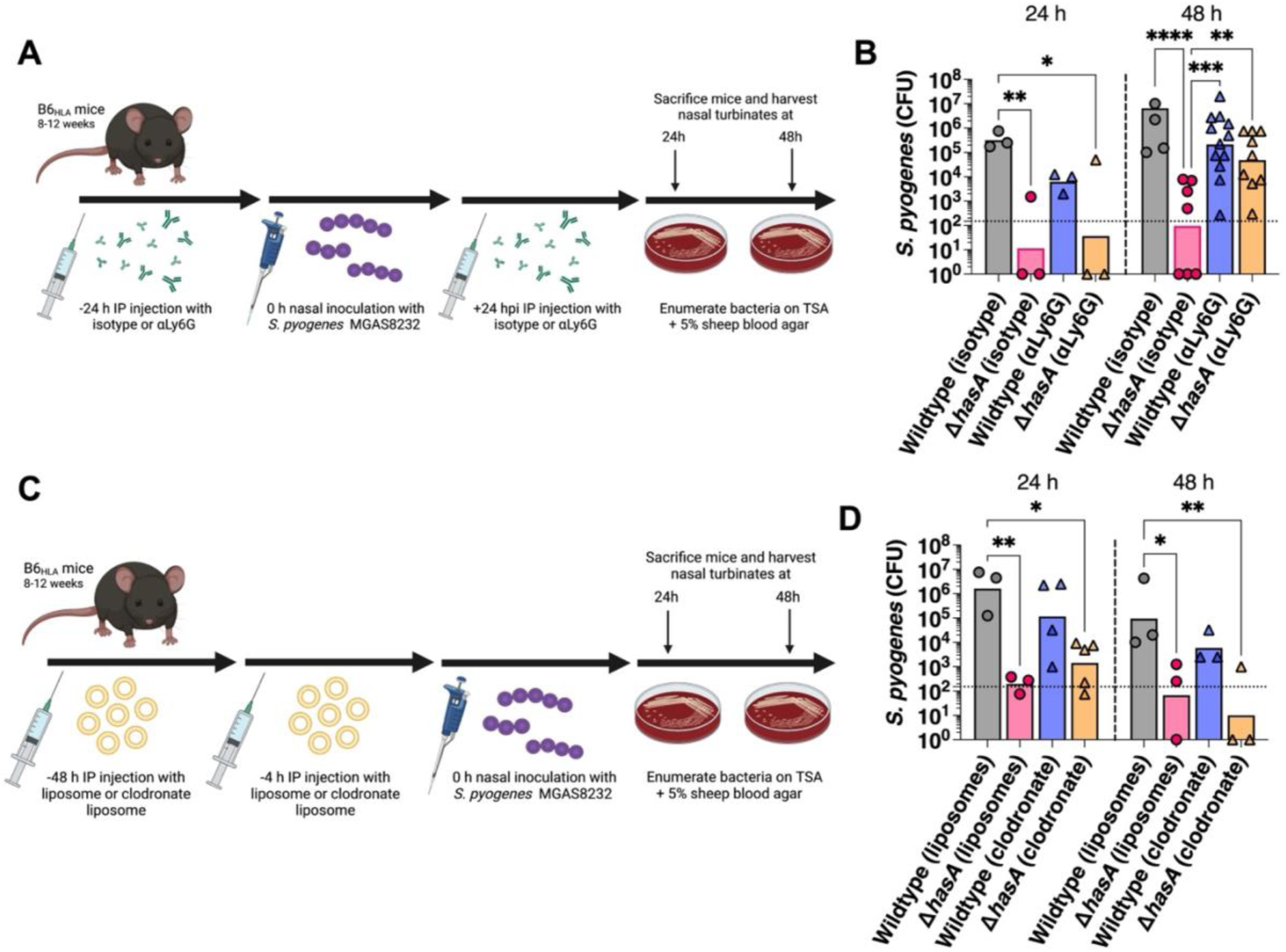
Early clearance of the HA capsule-deficient mutant from murine nasal turbinates is due to enhanced susceptibility to neutrophil-mediated killing. (A) Schematic outline for *in vivo* depletion of neutrophils with injections of 250 µg (500 µg total) of αLy6G or isotype control rat IgG2a 24 h prior to and 24 h post-intranasal challenge with 10^8^ CFUs of *S. pyogenes* wildtype or Δ*hasA* mutant strains. (B) Neutrophil effects on *S. pyogenes* survival in the nasopharynx. Data points represent CFUs from cNTs of individual mice 24 and 48 h post-infection. Horizontal bars represent the geometic mean. The horizontal dotted line indicates limit of detection. Significance was determined by two-way ANOVA with Tukey’s multiple comparisons (*, *P* < 0.05; **, *P* < 0.01; ****, *P* < 0.0001). (C) Schematic outlining the clodronate liposome-based depletion of macrophages in B6_HLA_ mice prior to nasopharyngeal infection with ∼10^8^ CFUs of wildtype and Δ*hasA S. pyogenes* MGAS8232. (D) Bacterial burden in murine cNTs at 24 and 48 h post-infection are shown. Each point represents *S. pyogenes* CFUs from individual mice, and the horizontal bars represent the geometric mean. Significant differences were determined by two-way ANOVA with Tukey’s multiple comparisons (*, *P* < 0.05; **, *P* < 0.01).

We next wanted to determine if preventing immune evasion by the HA capsule was specific to neutrophils or if it was shared by other phagocytes. For this purpose, mice were depleted of macrophages by administering clodronate containing liposomes, or empty control liposomes, and infected intranasally with *S. pyogenes* (**Figure 3C**). Following treatment with clodronate liposomes, similar amounts of wildtype *S. pyogenes* were recovered from infected cNTs compared to control liposome-administered mice at both 24-and 48-hours (24h, *p* = 0.6826; 48h, *p* = 0.8534), indicating that macrophage depletion did not impact nasopharyngeal infection by wildtype *S. pyogenes* (**Figure 3D**). As expected, control liposome-administered mice showed a reduction of the Δ*hasA* mutant at both 24 hours (*p* < 0.01) and 48 hours (*p* < 0.05) compared to wildtype-infected control mice (**Figure 3D**). At 24-hours, there was a trend of ∼1 log-fold greater recovery of Δ*hasA* CFUs in macrophage-depleted mice; however, this was not statistically greater than Δ*hasA* control mice (*p* = 0.7883) and remained significantly lower than control wildtype-infected mice (*p* < 0.05) (**Figure 3D**). At 48-hours, CFUs from the Δ*hasA* mutant in macrophage-depleted mice remained significantly lower than control wildtype-infected mice (*p* < 0.01). Overall, macrophage depletion by clodronate liposomes did not rescue the low infection phenotype by the Δ*hasA* strain at either 24-or 48-hours, and did not significantly influence infection by wildtype *S. pyogenes.* These results suggest that protection against phagocytosis by macrophages is not a key mechanism by which the HA capsule functions during experimental nasopharyngeal infection.

### Bacterial burden and lesion pathology during skin infection are enhanced by *S. pyogenes* hyaluronic acid capsule expression

Pharyngeal colonization by *S. pyogenes* is believed to be the major reservoir for this pathogen in developed countries, yet skin infections (impetigo) tend to be more prevalent in resource-poor settings [48]. Since nasopharyngeal infection by *S. pyogenes* MGAS8232 was notably compromised by the loss of the HA capsule, we next performed a novel skin infection model to further assess whether capsule expression could promote experimental skin infection. To address this, B6_HLA_ mice were intradermally injected in each hind flank with 2.5×10^7^ CFUs of wildtype, Δ*hasA,* or Δ*hasA* + *hasA* strains. There was a ∼10% decline in the weights of mice infected with wildtype *S. pyogenes,* a striking contrast to mice infected with the Δ*hasA* strain that gained weight over the 72-hour infection period (**Figure 4A**). The Δ*hasA* mutant strain revealed a clear reduction in virulence through considerably smaller lesions and less inflamed tissue over the infection period compared to wildtype*-*infected mice (**Figures 4B and 4D**). Significantly less bacterial CFUs were also recovered from each Δ*hasA-*infected lesion compared to wildtype-infected lesions (**Figure 4C**). Though weight loss and lesion sizes were only partially restored in Δ*hasA* + *hasA-*infected mice, bacterial CFUs recovered were fully complemented and did not differ from wildtype-infected mice (**Figures 4B-D**). The moderate restoration of weight loss and lesion sizes in Δ*hasA* + *hasA*-infected mice may be due to the incomplete complementation of the *hasA* gene expressed *in trans* using the pDC*erm* plasmid, as erythromycin was not used topically to maintain plasmid expression and replication over the 72-hour infection period. This is supported by the observation of some capsule deficient reversion colonies on plated skin homogenates (data not shown). Overall, these data demonstrate that the expression of the HA capsule in *S. pyogenes* supports virulence during acute skin infections in B6_HLA_ mice.

**Figure 4.**
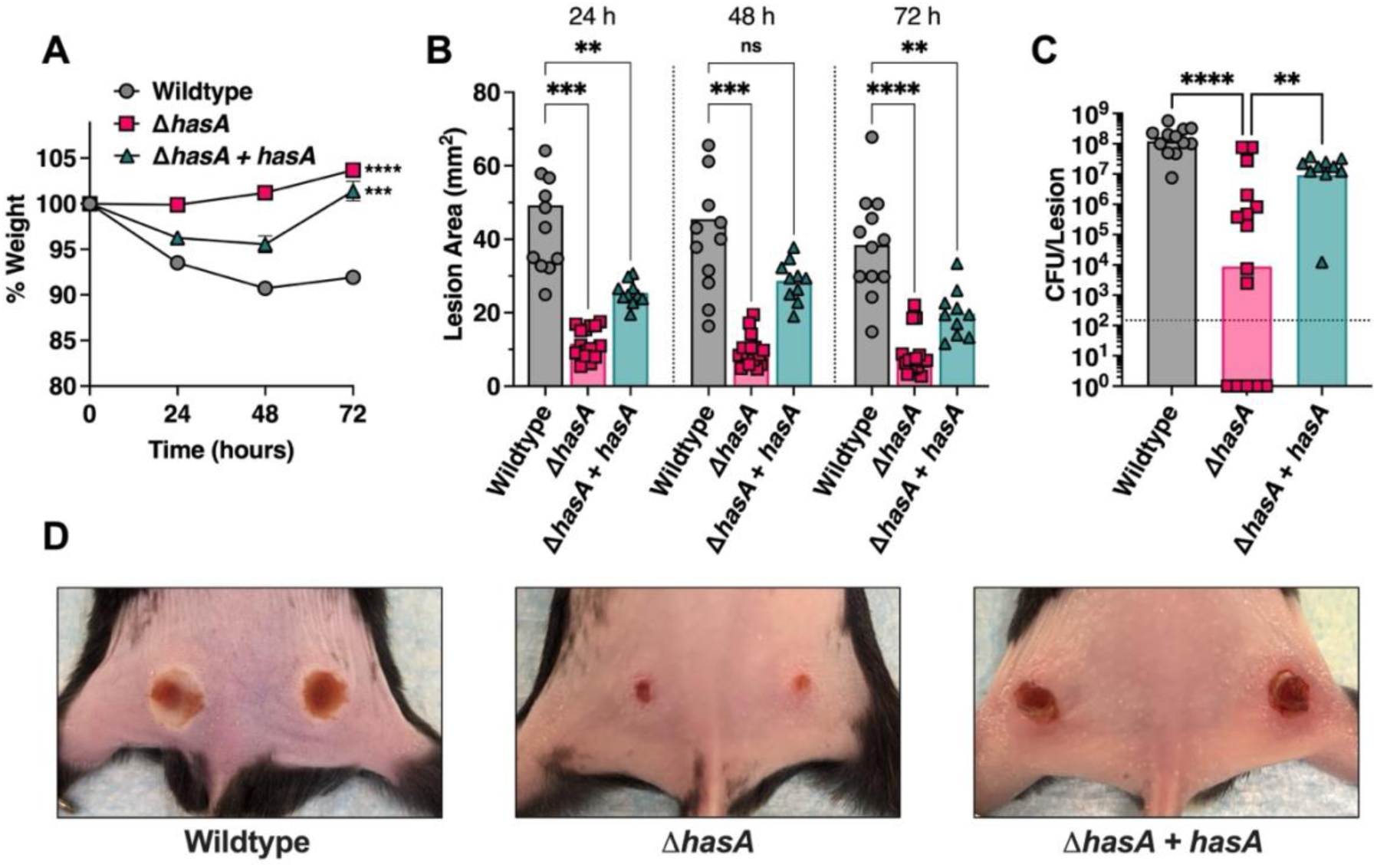
Hyaluronic acid capsule expression by *Streptococcus pyogenes* promotes skin infection in B6_HLA_ mice. B6_HLA_ mice were administered ∼5 × 10^7^ CFUs of wildtype *S. pyogenes* MGAS8232, or the *hasA* mutant, or the Δ*hasA + hasA* complemented strain by intradermal injections in each hind flank. **(A)** Weights of B6_HLA_ mice at 24-, 48-, and 72-h following skin challenge. Data is represented as a percentage of day 0 weight. Data points represent the weight means ± SEM. **(B)** Skin lesion areas of mice at 24-, 48-, and 72-h after skin challenge. Data points represent the mean. Significance was determined by two-way ANOVAwith Geisser’sGreenhouse correction and Dunnett’s multiple comparisons test (*, *P* < 0.05, **, *P* < 0.01, ***, *P* < 0.001;) for panels **A** and **B**. **(C)** Data points represent CFUs from individual infected skin abscesses per lesion from mice at 72 h. Horizontal bars represent the geometric mean. Significance was determined by one-way ANOVA with Kruskal-Wallis test (****, *P* < 0.0001; **, *P* < 0.01). The horizontal dotted line indicates the theoretical limit of detection. **(D)** Representative skin lesion images from B6_HLA_ mice 72 h following skin challenge.

### Depletion of neutrophils recovers *S. pyogenes* Δ*hasA* bacterial load during skin challenge

To examine whether a lack of neutrophils would similarly enhance infection by unencapsulated *S. pyogenes* in the skin, B6_HLA_ mice were depleted of neutrophils as described above and challenged with subdermal infections with 2.5×10^7^ CFUs wildtype or Δ*hasA S. pyogenes* MGAS8232. Irrespective of depletion status, mice infected with the Δ*hasA* mutant revealed significantly less weight loss and considerably smaller lesions compared to wildtype*-*infected mice over the infection period (**Figure 5A, 5B, and 5D**). Neutrophil depletion did not impact weight loss, lesion sizes, or the amount of wildtype *S. pyogenes* CFUs retrieved from each infected lesion (**Figure 5A-D**). As expected, control mice receiving the isotype antibody showed a significant reduction of Δ*hasA* mutants recovered within lesions at 72 hours (*p* < 0.001) compared to wildtype-infected control mice (**Figure 5C**). In contrast, neutrophil depleted mice displayed a sharp increase in Δ*hasA* CFUs recovered from each lesion compared to control mice that received infections with the Δ*hasA* mutant, despite presenting similar lesion sizes (**Figure 5C and 5D**). Overall, these data demonstrate that the *S. pyogenes* HA capsule is important for resisting bacterial killing by neutrophils during experimental skin infection.

**Figure 5.**
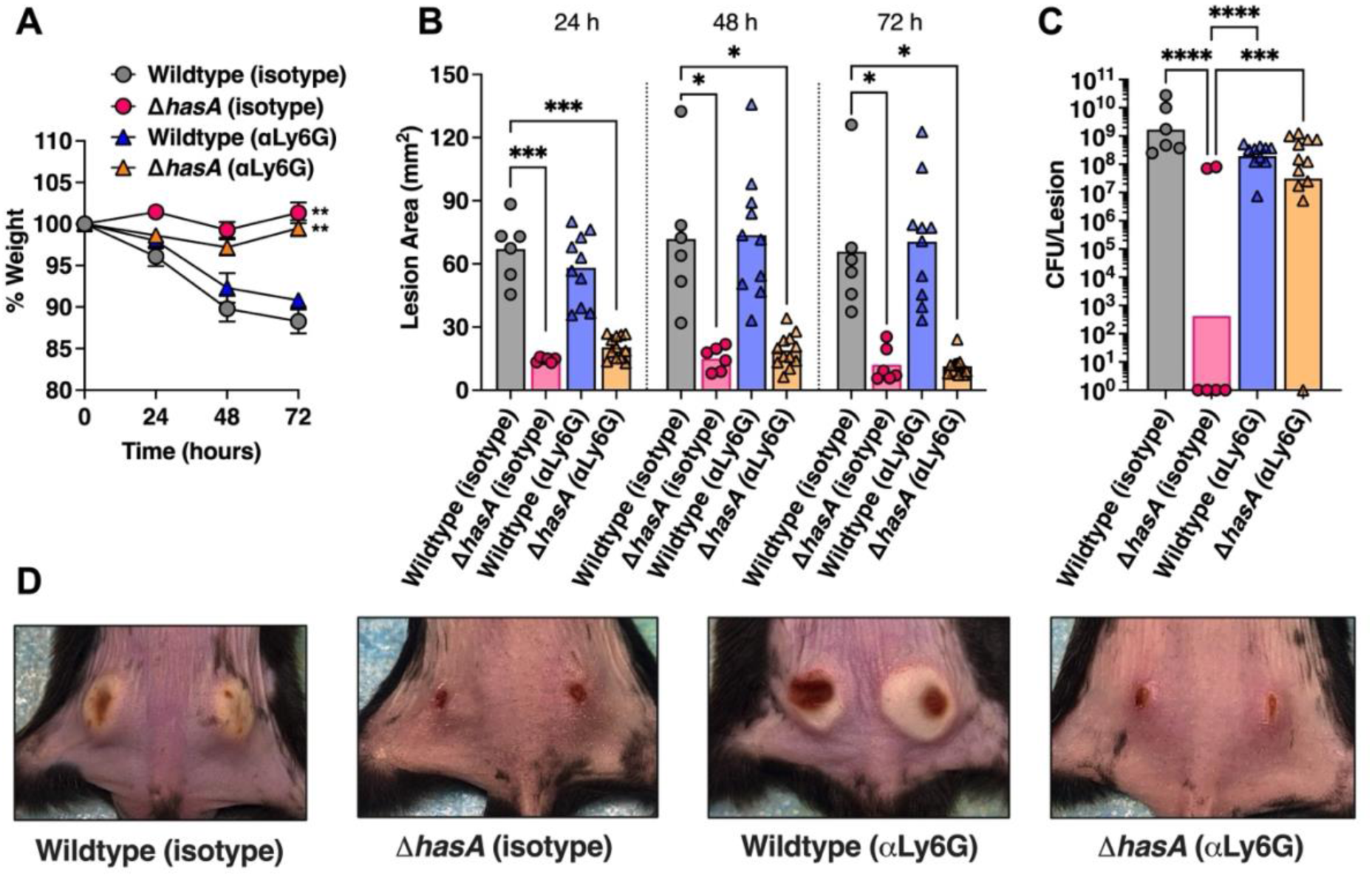
The hyaluronic acid capsule is important for resisting neutrophil-mediated killing in the skin. B6_HLA_ mice were administered ∼5 × 10^7^ CFUs of wildtype *S. pyogenes* MGAS8232 or the Δ*hasA* or Δ*hasA + hasA* complemented strain intradermally in each hind flank. Mice received αLy6G or rat IgG2a isotype control antibodies intraperitoneally 24-h preceding and 24-h after skin infections. **(A)** Weights of B6_HLA_ mice at 24-, 48-, and 72-h following *S. pyogenes* skin challenge. Data is represented as a percentage of day 0 weight. Data points represent the weight means ± SEM. **(B)** Skin lesion areas of mice following at 24, 48, and 72 h after skin challenge. Data points represent the mean. **(C)** Data points represent CFUs from individually infected skin abscesses per lesion from mice at 72 h. Horizontal bars represent the geometric mean. The horizontal dotted line indicates limit of detection. Significance was determined by two-way ANOVA with Geisser’s Greenhouse correction and Dunnett’s multiple comparisons test (*, *P* < 0.05, **, *P* < 0.01, ***, *P* < 0.001;) for panels **A** and **B**, or one-way ANOVA with Kruskal Wallis (***, *P* < 0.001; ****, *P* < 0.0001) for panel C. **(D)** Representative skin lesion images from B6_HLA_ mice 72 h following skin challenge.

## DISCUSSION

*S. pyogenes* is a human-specific bacterial pathogen and we previously demonstrated that mice that express human MHC-II molecules (B6_HLA_ mice) were dramatically more susceptible to experimental nasopharyngeal infection, denoting MHC-II as an important host factor for the adaptation of *S. pyogenes* to the human host [36]. It was further demonstrated that host sensitivity to superantigen-mediated T cell activation induces an excessive inflammatory signature within the nasopharyngeal environment that promoted the infection [49] and additionally, IL-1β-mediated inflammation mediated by the SpeB protease can similarly promote *S. pyogenes* colonization of the nasopharynx [47]. Consequently, *S. pyogenes* must remodel its external environment and balance superantigen-and SpeB-mediated inflammation while tempering host immune clearance mechanisms at various stages of infection, each of which may be influenced by strain-specific differences and tissue-specific cues that can affect the outcome of infection. Herein, we leveraged the B6_HLA_ mouse model to investigate the role of the HA capsule of *S. pyogenes* MGAS8232 during acute infections and provide evidence that the *S. pyogenes* capsule functions *in vivo* to inhibit neutrophil-mediated clearance in both experimental nasopharyngeal and skin infections.

The findings presented here illustrate an important role of the HA capsule during the pathogenesis of acute upper respiratory and skin infections by *S. pyogenes*; however, this may appear inconsistent with some previous investigations for other encapsulated bacterial pathogens. For example, reduced or eliminated capsule production appears to have advantages for the invasive potential or persistence at mucosal surfaces across multiple bacterial species, including *Streptococcus agalactiae* (Group B *Streptococcus*) [50]*, Streptococcus pneumoniae* [51]*, Neisseria meningitidis* [52, 53], and *Haemophilus influenzae* [54]. *S. pneumoniae*, for example, varies capsule expression from its initial abundance to prevent mucus-mediated clearance [55], yet it is subsequently downregulated to expose underlying adherence molecules [56] and to promote biofilm formation [57, 58]. Although the Δ*hasA* mutant did have enhanced invasion of epithelial cells (**Figure 1D, 2D**), we did not detect an increase in the dissemination to other organs *in vivo* (**Figure S1**). Furthermore, our findings also contradict some previous reports where acapsular *S. pyogenes* infected the pharynx as effectively as the parental strain in a baboon model of pharyngeal infection [18], and that frameshift inactivating mutations in the *hasA* or *hasB* genes that deplete capsule production contributed to persistence during asymptomatic carriage [18, 29]. The use of different strains, and different infections models, have likely have contributed to these disparate findings, and since not all *S. pyogenes* strains encode the *has* operon [59], it is clear that the HA capsule is not an essential virulence factor for all *S. pyogenes* isolates. Nevertheless, our work is entirely consistent with other prior work demonstrating an important selective advantage of the HA capsule for survival within the nasopharynx [15,17,18,24].

During infection of the nasopharynx, the epithelium and mucus layer form the frontline barrier against invading pathogens where adherence to epithelial cells or exposed ECM may be exploited to prevent mucosal-mediated removal. In this study, we report that unencapsulated *S. pyogenes* display reduced binding to collagen type IV and fibronectin ECM components (**Figure 2A and 2B)**, yet pharyngeal epithelial cell adhesion and internalization were significantly greater compared to the encapsulated wildtype strain (**Figure 2C and 2D**). Although this may appear to be paradoxical, prior studies have shown that once internalized within epithelial cells, *S. pyogenes* is rapidly killed [24]. Thus, entry into cells is unlikely a virulence mechanism, but rather a failure of *S. pyogenes* to avoid ingestion by host cells. Therefore, encapsulation helps resist internalization and enhances the capacity to invade tissues by an extracellular route to promote *S. pyogenes* infection. Future work is needed to clarify adherence properties of encapsulated and unencapsulated *S. pyogenes,* however, an impairment in adherence appears to be less important throughout the overall course of infection compared to the capsule’s potent protective effect from ingestion and killing by host phagocytes.

Upon infection with *S. pyogenes*, the immune system launches a complex innate response that largely depends on the recruitment and activity of neutrophils, macrophages, and dendritic cells [60–64]. In this study we have demonstrated the the HA capsule is a key structure produced by *S. pyogenes* that promotes resistance to neutrophil mediated killing. Neutrophils are the most abundant leukocyte involved in innate host responses, acting as both professional detectors that release inflammatory alarms to invading bacteria as well as direct killers via phagocytosis, degranulation, and the formation of neutrophil extracellular traps (NETs). While neutrophil influx during severe infections is protective against *S. pyogenes* [63], we show that depleting neutrophils or macrophages did not affect wildtype *S. pyogenes* MGAS8232 acute infections. These results are in contrast to findings where neutrophils are key for pathogenesis and that neutrophil ablation by αLy6G administration reduces *S. pyogenes* infection of the nasopharynx [47, 65]. However, conventional C57BL/6 mice were used in these studies with superantigen-mediated inflammation absent. In the presence of a superantigen-driven inflammatory response capable of promoting infection [36], our results indicate that neutrophils are not essential for *S. pyogenes* to establish nasopharyngeal or skin infections. Instead, expression of the HA capsule offered a clear survival advantage that promoted a strong resistance to bacterial clearance by neutrophils. Since innate immune cells are thought to participate in host protection against *S. pyogenes,* more research is needed to define specific roles, to examine crosstalk, and to address redundancy in responses between individual cell types.

Interestingly, both encapsulated and unencapsulated type 18 *S. pyogenes* are equally opsonized by C3 in either plasma or serum [25], suggesting that the HA capsule does not inhibit complement activation or deposition of complement fragments on the bacterial cell wall. Since opsonization does not necessarily lead to phagocytic ingestion, the HA capsule may serve as a physical barrier that interferes with leukocyte access to opsonic complement proteins deposited on the bacterial surface [25]. More recently, the HA capsule has been shown to promote bacterial survival within NETs by resisting a major component and antimicrobial effector, cathelicidin antimicrobial peptide LL-37 [66]. As different strains of *S. pyogenes* harbour variations in global virulence factor expression, and consequently, express varying amounts of HA capsule, it is likely that distinct strategies to prevent phagocytic ingestion and killing are exploited among individual strains. For example, mutations that produce a truncated RocA (<u>r</u>egulator <u>o</u>f <u>C</u>ov) protein have amplified expression of the *has* operon through transcriptional activation of the repressor *covR*, and have been identified in *S. pyogenes* types *emm*18 and *emm*3 [34, 67]. Thus, no single strain of *S. pyogenes* should be considered representative of the population as a whole and future studies using additional encapsulated strains are recommended to draw general conclusions on the mechanisms utilized by the HA capsule. Although various strains may vary greatly in their degree of encapsulation, the results presented here provide evidence that HA capsule expression by *S. pyogenes* MGAS8232 promotes a strong resistance to killing by neutrophils during acute infection models. Defining strategies by which neutrophils can counteract HA capsule resistance is warranted to combat this leading bacterial pathogen.

## ACKNOWLEDGEMENTS

This work was supported by an operating grant from the Canadian Institutes of Health Research (CIHR) to JKM. JRH was supported in part by RGE Murray Graduate Scholarship and Queen Elizabeth II Graduate Scholarship in Science and Technology. BAS was supported in part by Ontario Graduate Scholarship. Figure 3A and 3C were created using Biorender.com.

## AUTHOR CONTRIBUTIONS

JRH executed the majority of the experimental work with assistance by BAS, SWT and HCC. Experimental design and data interpretation were performed by JRH with assistance from SWT. JRH and JKM conceptualized the study and wrote the manuscript, which was reviewed and approved by all co-authors.

## DECLARATION OF INTERESTS

The authors have declared that no competing interests exist.

## MATERIALS & METHODS

### Bacterial strains, media, and growth conditions

Bacterial strains used in this study are listed in **Table 1**. The main bacterial model strain for our work is *S. pyogenes* MGAS8232, an M18 serotype and pharyngeal isolate from a patient with acute rheumatic fever [33]. *S. pyogenes* strains were grown statically in Todd Hewitt broth (BD Biosciences; Franklin Lakes, NJ, USA) supplemented with 1% (w/v) yeast extract (BD Biosciences) (THY) and 1 μg mL^-1^ erythromycin when appropriate. For solid media preparation, 1.5% agar and/or 1 μg mL^-1^ erythromycin were added to the media when applicable. Molecular cloning experiments utilized the *E. coli* XL1-Blue strain cultured in Luria-Bertani (LB) broth (Thermo Fisher Scientific, Waltham, MA, USA) aerobically at 37°C, or Brain Heart Infusion (BHI; BD Biosciences, Franklin Lakes, NJ, USA) media containing 1.5% (w/v) agar (Thermo Fisher Scientific). Media was supplemented with 150 μg mL^-1^ erythromycin (Sigma-Aldrich Canada, Oakville, ON, Canada) as necessary. A complete list of plasmids used in this study can be found in **Table 1**.

### Construction of recombinant *S. pyogenes* strains

References to genomic loci are based on the genome of *S. pyogenes* MGAS8232 [33]. In-frame genetic deletion in the *hasA* gene was generated using the Gram-positive-*E. coli* shuttle vector, pG^+^host5 [32] (**Table 1**). Using appropriate PCR amplification primers listed in **Table 2**, the upstream and downstream regions flanking the *hasA* gene were amplified from the MGAS8232 genome. Allelic replacement of the wildtype *hasA* gene with the deletion mutant Δ*hasA* via homologous recombination was conducted as described previously [36]. For complementation of the *hasA* genetic deletion, DNA fragments containing the *hasA* open reading frame and its native promoter were amplified from the *S. pyogenes* MGAS8232 genome using complementation primers listed in **Table 2**, and cloned into the XhoI and SpeI restriction sites of the plasmid pDC*erm* [35]. This construct (pDC*erm*::*hasA*) was electroporated into the MGAS8232 Δ*hasA* mutant strain to produce the *hasA* complementation strain (Δ*hasA + hasA*).

### Genomic sequencing analysis

Genomic DNA preparations from *S. pyogenes* MGAS8232 wildtype and Δ*hasA* strains were sent for paired end Illumina sequencing at the John P. Robarts Research Institute sequencing facility (University of Western Ontario, London, Ontario). Illumina short-read sequence data were used to generate *de novo* assemblies using SPAdes v3.15 [68], which were annotated using Prokka v1.12 [69]. These assemblies have been deposited at NCBI. Any sequence differences between the strains were determined using Snippy v4.1 (https://github.com/tseemann/snippy). The publicly available *S. pyogenes* MGAS8232 sequence was used as a reference (Bioproject: PRJNA286). SNPs unique to the Δ*hasA* mutant were reported (**Table S2**).

### Extracellular matrix binding assay

Corning Costar 9018 high-binding 96-well plates (Corning; Kennebuck, ME, USA) were coated with 1 μg of collagen type IV (Sigma-Aldrich) or fibronectin (Calbiochem, EMD Millipore Corporation; Temecula, CA, USA) dissolved in carbonate-bicarbonate buffer (0.2M sodium carbonate anhydrous, 0.2M sodium bicarbonate, pH = 9.6) and left overnight at 4°C. The following day, the plates were washed three times using PBS with 0.05% (v/v) tween-20 and blocked for two hours with 5% (w/v) skim milk at room temperature and then washed as described. Afterwards, 100 μlof bacteriacontaining 10^7^ CFUs were added in triplicate to pre-coated wells and left for 2.5 hours at 37°C. Plates were washed and fixed with 10% neutral buffered formalin (VWR International; Randor PA, USA) for 40 minutes and then washed again. Wells were incubated with 50 μlof 0.5% (w/v) crystal violet(Sigma-Aldrich) in 80% (v/v) sterile MilliQ water and 20% (v/v) methanol for 5 minutes at room temperature before being washed five times. Stain was solubilized in 5% (v/v) acetic acid with mild agitation for 10 minutes. Colorimetric analysis was measured at OD590 using Synergy HTX Multi-Mode Microplate Reader (Biotek).

### Human cell culturing

The Detroit-562 (ATCC CCL-138) human pharyngeal cell line was grown and maintained at 37°C in 5% CO_2_ in minimal essential medium (MEM) and passaged every 2-3 days. All tissue culture basal media was supplemented with 10% (v/v) heat-inactivated fetal bovine serum (FBS; Sigma-Aldrich, St. Louis, MO, USA), 100 IU penicillin (Gibco®, Life Technologies Inc, Carlsbad, CA, USA), and 100 μg mL^-1^ streptomycin (Gibco-BRL, Life Technologies, Grand Island, N.Y.), all filtered through a 0.2 μm PES filter (Nalgene^TM^, Thermo Scientific, Waltham, MA, USA).

### Epithelial cell adhesion and invasion

Detroit-562 cells were grown to confluence on 12 or 24-well TC-treated plates (Falcon, Corning). On the day of infection, cells were washed with PBS and maintained in their respective serum-free media for at least 1 h prior to infection. The average number of cells per well was calculated and used to determine the number of bacteria required for a multiplicity of infection (MOI) of 100. During this time, overnight streptococcal cultures were subcultured into pre-warmed media and grown to early exponential phase. Tissue culture plate wells were washed three times with PBS and inoculated with 500 μlof theresuspended bacterialdoseper welland left for 2-3 hours at 37°C in a 5% CO_2_ incubator to allow bacteria to adhere to the cells. Background adherence levels were measured by inoculating bacteria onto uncoated wells. Following incubation, wells were washed three times with PBS to remove non-adherent bacteria, and cell monolayers were lysed with 500 μl of 0.01% Triton X-100 for 5 minutes, followed by disruption of the wells by scraping with a 1 mL pipette tip. Solubilized wells were serially diluted 10-fold and plated onto TSA 5% sheep blood agar plates (BD Biosciences) to enumerate bacteria present.

For invasion experiments, bacterial were inoculated onto cell monolayers for 2-3 hours at 37°C with 5% CO_2_ and washed as above, followed by the addition of media containing 100 μg mL^-1^ of gentamicin for 1 hour at 37°C and 5% CO_2_ to kill extracellular bacteria. Wells were then extensively washed to remove the gentamicin media. Cells were lysed, serially diluted, and plated as described above to estimate the number of intracellular bacteria.

### Human ethics statement

Human venous blood was taken from healthy volunteer donors in accordance with human subject protocol 110859. The full study protocol was approved by the London Health Sciences Centre Research Ethics Board (University of Western Ontario, London, ON, Canada). Volunteers were recruited by a passive advertising campaign within the Department of Microbiology and Immunology at the University of Western Ontario, and following an outline of the risks, written consent was obtained from each volunteer before samples were taken. Following blood collection, samples were fully anonymized and no information regarding the identity of the donor, including sex and age, were retained.

### Whole blood survival assay

Lancefield bactericidal assays were performed as previously described [70] with some modifications. Briefly, a volume of 10 μl containing 1000 bacterial CFUs was added to 990 μl heparinized whole human blood in 1.5 mL Eppendorf tubes and allowed to incubate at 37°C with rotation. After 30, 60, 90, 120, and 180 minutes, samples were serially diluted 10-fold and drop plated in triplicate onto 5% TSA blood agar plates to enumerate the surviving bacteria in whole blood at indicated time points.

### Isolation of human polymorphonuclear neutrophils

Polymorphonuclear neutrophils (PMNs) were isolated using a Ficoll (GE-Healthcare) Histopaque (Sigma-Aldrich) density gradient centrifugation method. Briefly, heparinized blood from human donors was diluted with an equal volume of PBS (Wisent Bioproducts Inc.) and layered carefully onto a dual Ficoll-Histopaque gradient and centrifuged at 396 × g for 20 minutes without braking. The PMN layer was collected and washed with cold RPMI containing 0.05% human serum albumin (RPMI-HSA) and with addition of 1 mL ice-cold water to lyse residual erythrocytes. PMNs were then collected in RPMI-HSA following centrifugation and adjusted to 4 × 10^6^ cells mL^-1^.

### PMN survival assay

Overnight streptococcal cultures were grown to early exponential phase and diluted to 10^4^ CFU mL^-1^ in RPMI containing 10% (v/v) normal serum for 30 minutes to assist with bacterial opsonization. A volume of 0.225 mL opsonized bacteria were co-cultured with 0.025 mL of isolated human PMNs at 4 × 10^6^ cells mL^-1^ (1:10 bacterial CFUs to neutrophils) at 37°C with vigorous shaking. Control samples had no PMNs added to control for bacterial growth. Viable bacteria in each reaction mixture were measured after 60 mins by lysing cells with 750 µl of 0.025% Triton X100, followed by serial dilution and plating onto 5% TSA blood agar plates overnight at 37°C. Percent bacterial survival was calculated as average bacterial CFUs in the presence of neutrophils divided by bacterial CFUs in no PMN control samples.

### Mice

All mouse experiments were conducted in accordance with the Canadian Council on Animal Care Guide to the Care and Use of Experimental Animals. The Animal Use Protocol (AUP) number 2020-041 was approved by the Animal Use Subcommittee at the University of Western Ontario (London, ON, Canada). These mice were bred in a barrier facility at the University of Western Ontario and genotyped routinely for appropriate transgene expression. Human MHC class II transgenic (B6_HLA_) mice [71] were bred from McCormick laboratory colonies specifically for this study and were on a C57BL/6 (B6) background. During all breeding and experiments, mice were provided food and water *ad libitum* and appropriate enrichment was provided in all cages.

### Nasopharyngeal infection model

Preparation of *S. pyogenes* MGAS8232 for nasal inoculation has been previously described [36,37,49]. Briefly, bacteria were grown to early exponential phase (OD_600_ of 0.2–0.3), cells were centrifuged, washed, and resuspended in Hank’sbuffered salinesolution (HBSS) at∼1 × 10^8^ CFUs per 15 μL. Mice were given 2 mg ml^-1^ neomycin sulphate *ad libitum* in their drinking water two days prior to infection to reduce the nasal microbiota. Mice were anesthetized using FORANE (isoflurane, USP;Baxter Corporation;Mississauga, ON, Canada) and 7.5 μLof bacterialinoculum was administered into each nostril. Mice were sacrificed 24-or 48-h post-infection and their complete nasal turbinates (cNTs), including the nasal-associated lymphoid tissue, nasal turbines, and maxillary sinuses, were extracted and homogenized using a glass homogenizer [37]. Murine lungs, liver, spleen, kidneys, and heart were also removed for bacterial enumeration where indicated. Organswereserially diluted in HBSSand plated onTSAsupplemented with 5% sheep’s blood for bacterial enumeration. Counts less than 30 CFUs per 100 μL of cNT were considered below the theoretical limit of detection.

### Skin infection model

The fur on the lower backs of B6_HLA_ mice between 8-12 weeks old was removed by shaving and hair removal cream the day prior to infection. *S. pyogenes* was grown and prepared as stated above and resuspended to 5 × 10^8^ CFU per mL in HBSS. Mice were anesthetized and a 50 μL dose containing ∼2.5 × 10^7^ CFU was injected intradermally into each lower flank. On each day following infection, mice were weighed and lesions at the injection sites were measured using calipers. At 72-hours post-infection, mice were sacrificed and the skin around each injection site was harvested, homogenized, and plated on TSA with 5% sheep’s blood overnight at 37°C for bacterial enumeration. Bacterial burden was presented as CFUs from individual lesions.

### Immunofluorescent histology

At the previously identified endpoint, cNTs were collected as described [37] and tissues were fixed in periodate-lysine-paraformaldehyde (PLP) and prepared for sectioning as previously [72]. Following fixation, cNT were passed through sucrose gradients and frozen in OCT (TissueTek) media. Serial sections (7 µm) were cut using a cryostat. Prior to staining, all slide-mounted tissue sections were blocked with PBS containing 1% BSA, 0.1% Tween-20, and 10% rat serum. Sections were stained with the following antibodies: anti-*Streptococcus pyogenes* Group A Carbohydrate (Abcam, ab9191), anti-B220 (Biolegend, RA3-6B2), anti-CD3 (Biolegend, 17A2), and anti-Ly6G (Biolegend, 1A8). After staining, sections were mounted with ProLong Gold Antifade Reagent (Invitrogen). Tiled images of whole cNT sections were collected using a DM5500B fluorescence microscope (Leica) at 10× and 20×.

### PMN and macrophage depletion *in vivo*

The function of neutrophils and macrophages during acute infections by *S. pyogenes* were examined in B6_HLA_ mice between 8-12 weeks old. Neutrophils were depleted *in vivo* by intraperitoneally injecting mice with 250 μg mAb αLy6G clone 1A8 (BioXcell, NH, USA) 24 h before and 24 h following nasopharyngeal and skin infections. Control mice received rat IgG2a clone 2A3 (BioXCell, NH, USA). Depletion of circulating neutrophils has been confirmed in previous studies using flow cytometry of Ly6G^+^ expressing populations in blood [47,73,74]. The depletion of macrophages in B6_HLA_ mice between 8-12 weeks old has similarly been previously described [75–78]. Briefly, 200 μL Clodronate containing liposomes and control liposomes [Clodrosome® + Encapsome® (Encapsula Nano Sciences)] were intraperitonially injected in mice at both 48-and 4-h prior to nasopharyngeal infection with *S. pyogenes.* Bacterial burden in nasal turbinates was examined at both 24-and 48-h following infection.

### Detection of cytokines and chemokines *in vivo*

Cytokine and chemokine concentrations were determined from cNT homogenates of B6_HLA_ mice infected with *S. pyogenes* MGAS8232. Uninfected murine cNT homogenates were measured as a background control. Mutiplex cytokine array (Mouse Cytokine/Chemokine Array 32-Plex) was performed by Eve Technologies (Calgary, AB, Canada) and data on heat maps is presented as normalized median cytokine responses (X_normalized_ = [(x -x_min_)/(x_max_ – x_min_)]) from cNT homogenates.

### Statistical analysis

All statistical analysis was completed using GraphPad Prism 9.3.1. Significance was calculated using the Student’s *t* test or one-way ANOVA with Dunnett’s or Tukey’s multiple comparisons post hoc test where indicated. A *P* value less than 0.05 was determined to be statistically significant.

## SUPPLEMENTAL DATA

**Figure S1.**
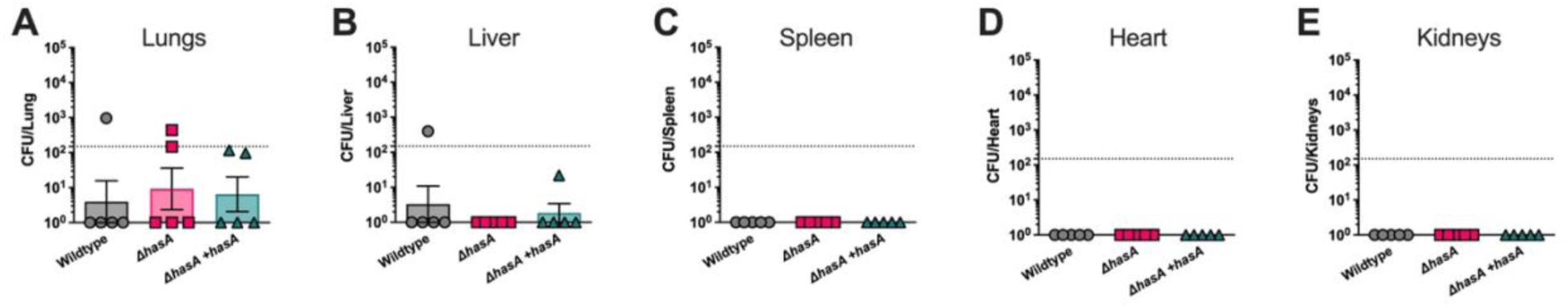
Deletion of the *hasA* gene in *S. pyogenes* MGAS8232 does not enhance bacterial dissemination in B6_HLA_mice. B6_HLA_mice were nasally challenged with ∼10^8^CFUs of *S. pyogenes* wildtype or Δ*hasA*. Mice were sacrificed 48-h later, and indicated organs were harvested, homogenized, and plated on TSA with 5% sheep blood agar to assess bacterial dissemination. Bacterial CFUs were measured in the **(A)** lungs, **(B)** liver, **(C)** spleen, **(D)** heart, and **(E)** kidneys. Data points represent CFUs of indicated organs from individual mice (n ≥ 4 per group). Bars represent mean ± SEM. Horizontal dotted line indicates theoretical limit of detection. Significance was determined by one-way ANOVA with Dunnett’s multiple comparisons test, data not significant.

**Figure S2.**
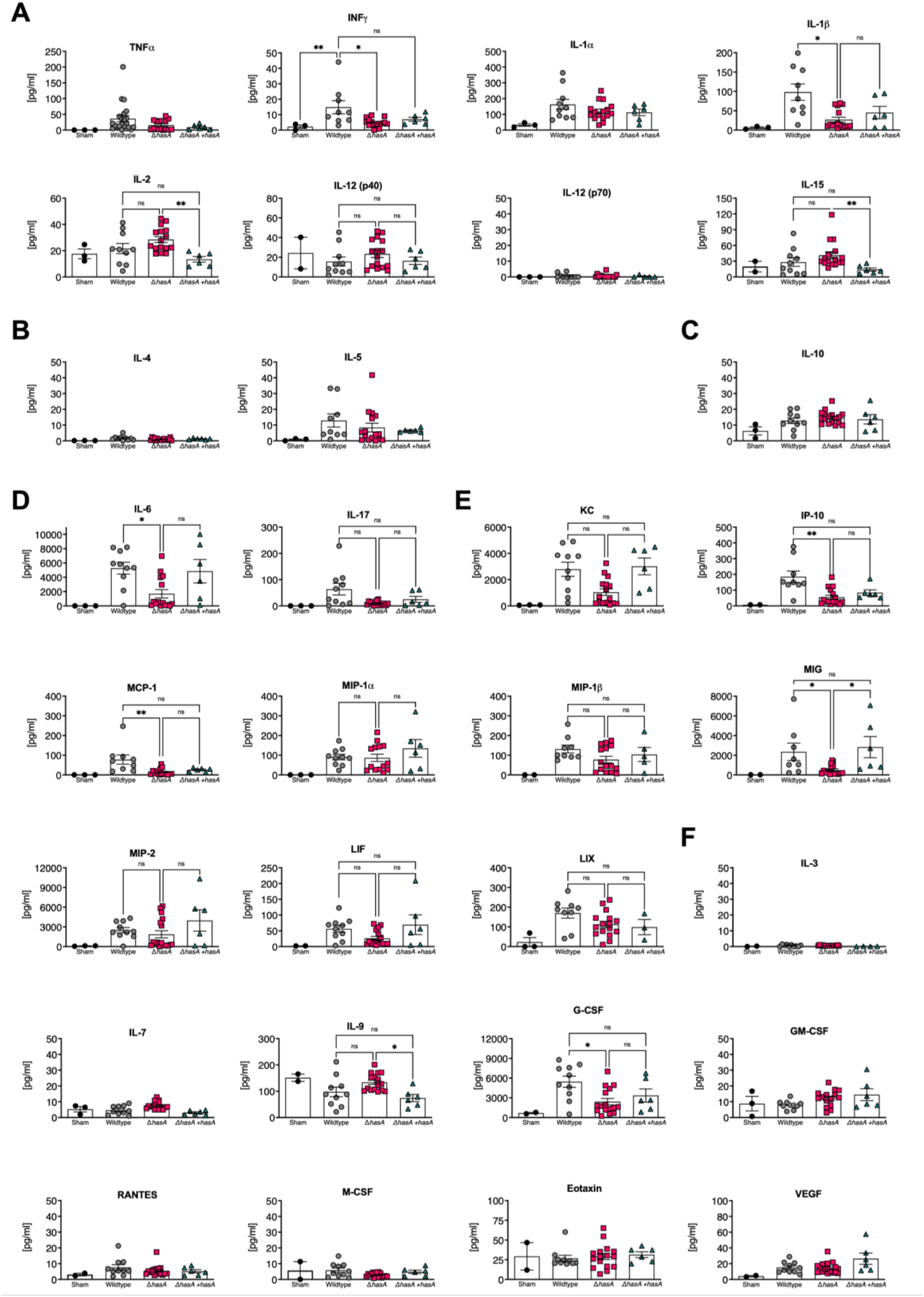
Cytokine response in nasal turbinates of B6_HLA_ mice during streptococcal infection. Mice were inoculated with HBSS as sham control (black circles) or infected intranasally with 10^8^ CFUs of wildtype *S. pyogenes* MGAS8232 wildtype, Δ*hasA,* or Δ*hasA* + *hasA* strains. Mice were sacrificed 48-h post-infection and cNT homogenates were analyzed for multiple cytokines and chemokines (Th1-type [**A**]; Th2-type cytokines [**B**]; Treg cytokines [**C**]; Th17 cytokines [**D**]; chemokines [**E**]; or growth factors [**F**]). Data represents the mean ± SEM of cNT cytokine/chemokine concentrations (pg mL^−1^) (*n* ≥ 3 mice per group). Significance was determined by one-way ANOVA with Dunnett’s multiple comparison test (* *P* < 0.05; ** *P* < 0.01; *** *P* < 0.001).

**Table S1.**
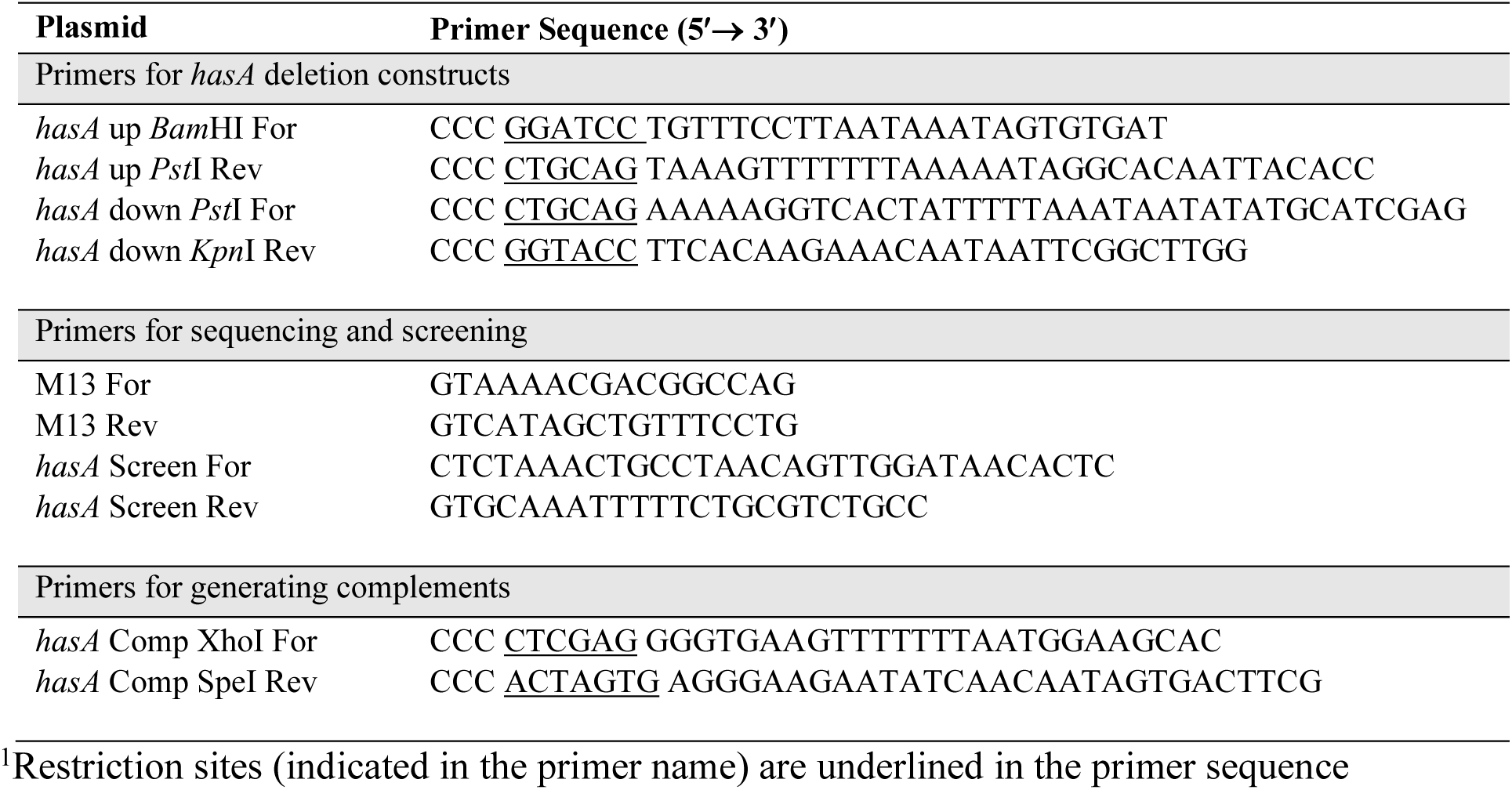
Primers in this study.

**Table S2.** SNPs identified through genome wide comparisons of *S. pyogenes* MGAS8232 wildtype and Δ*hasA* strains.

| Gene | Locus | Protein | Gene length (bp) | Codon number | Reference nucleotide | Competing nucleotide | Nucleotide in codon | Amino acid change |
| --- | --- | --- | --- | --- | --- | --- | --- | --- |
| pstI<sup>a</sup> | spyM18_1384 | AAL97979.1 | 1734 | 194 | A | T | 1 | L > F |
| pstI<sup>a</sup> | spyM18_1384 | AAL97979.1 | 1734 | 306 | C | T | 2 | S > F |

<sup>a</sup> cytosolic enzyme I in the phosphoenolpyruvate-protein phosphotransferase system

## Notes

### Competing Interest Statement

The authors have declared no competing interest.

